# Decynium-22 affects behavior in the zebrafish light/dark test

**DOI:** 10.1101/2021.01.14.426728

**Authors:** Caio Maximino

## Abstract

Decynium-22 (D-22) is an inhibitor of the uptake_2_ system of monoamine clearance, resulting in increased levels of dopamine, norepinephrine, and serotonin in the nervous system and elsewhere. Uptake_2_ is mediated by low-affinity, high-capacity transporters that are inhibited by glucocorticoids, suggesting a mechanism of fast glugocorticoid-monoamine interaction in the brain. D-22 dose-dependently increased anxiety-like behavior in adult zebrafish exposed to the light/dark test, monotonically increasing scototaxis (dark preference), but affecting risk assessment with an inverted-U-shaped response. These results suggest that the uptake_2_ system has a role in defensive behavior in zebrafish, presenting a novel mechanism by which stress and glucocorticoids could produce fast neurobehavioral adjustments in vertebrates. **Data:** https://github.com/lanec-unifesspa/decynium22

## 1. Introduction

Clearance of the monoamine neurotransmitters dopamine (DA), norepinephrine (NE), and serotonin (5-HT) released in the synaptic cleft is executed by two distinct mechanisms, uptake_1_ and uptake_2_ (Daws 2009a). Uptake_1_ is mediated by high-affinity, low-capacity transporters which include the NE transporter (SLC6A2, NET), the DA transporter (SLC6A3, DAT), and the serotonin transporter (SCL6A4, SERT)(Rudnick *et al*. 2014). This SLC6A family has been implicated in the pathophysiology of mental disorders, including alterations of anxiety (Mohammad *et al*. 2016; Sullivan *et al*. 1999; Maximino 2012), and are the target for major classes of anxiolytic drugs, including tricyclic antidepressants, selective 5-HT reuptake inhibitors, and 5-HT-NE reuptake inhibitors (Gorman 2002). Uptake_2_ is mediated by low-affinity, high-capacity transporters which include organic cation transporters (OCT1-3; SLC22A1-3) and the plasma membrane monoamine transporter (PMAT; SLC29A4) (Wu *et al*. 1998; Zhou *et al*. 2007). Evidence has suggest that uptake_2_ plays significant roles in the regulation of monoaminergic neurotransmission and maintenance of homeostasis (Hagan *et al*. 2011; Zhu *et al*. 2012; Rahman *et al*. 2008; Daws *et al*. 2013; Wultsch *et al*. 2009; Horton *et al*. 2013; Kitaichi *et al*. 2005; Schildkraut and Mooney 2004). Uptake2 is an interesting system because not only it is a system best suited for extraneuronal uptake (due to its low-affinity, high-capacity, “promiscuous” characteristic), but also because it is blocked by glucocorticoids (Hill *et al*. 2011; Wu *et al*. 1998). As a result, uptake_2_ represents an intersection in the pathophysiology of stress and anxiety, a mechanism by which circulating glucocorticoids (GCs) can rapidly increase monoamine levels in the brain (Maximino 2012). Uptake_2_ has been shown to participate in anxiety-like behavior: SLC22A3 knockout mice show decreased anxietylike behavior in the open field test and in the elevated plus-maze (Wultsch, Grimberg, Schmitt, Painsipp, Wetzstein, Frauke, et al., 2009; but see Vialou et al., 2008). Knockdown of SLC22A3 expression in the brains of mice decreases immobility time in the forced swimming test (Kitaichi *et al*. 2005), a screen for antidepressant-like effects (Willner 1984). Finally, while decynium-22 (D-22), an uptake_2_ inhibitor, had no behavioral effect by itself, co-treatment with fluvoxamine produced synergistic effects on 5-HT clearance and immobility in the forced swimming test (Horton *et al*. 2013).

Zebrafish *(Danio rerio* Hamilton 1822) have been proposed as model organisms in the study of behavioral functions and its disorders (Stewart *et al*. 2015; Gerlai 2014; Maximino *et al*. 2010b; Rinkwitz *et al*. 2011). The advantages of using this species in behavioral studies stem from its use in developmental biology (i.e., small size, fast generation times, high reproduction rates) and the availability of tools to image and manipulate its nervous system (Rinkwitz *et al*. 2011). Zebrafish demonstrate a robust endocrine response to acute stressors (Idalencio *et al*. 2017b; Idalencio *et al*. 2015; Tran *et al*. 2014; Fuzzen *et al*. 2010); importantly, simple acute stressors such as net chasing induce robust behavioral responses which are blocked by 5-HT reuptake inhibitors (Giacomini *et al*. 2016; Abreu *et al*. 2017) and DAergic and NErgic drugs (Idalencio *et al*. 2017a; Idalencio *et al*. 2015).

Currently, it is unknown whether zebrafish possess a functional uptake_2_ system. 14 *slc22* genes have been identified in zebrafish, and OCT3 appears absent (Popović 2014); *oct1* shows moderate expression in the brain, suggesting a role in neurotransmitter homeostasis (Popović 2014). No current information exists for PMAT as well. Nonetheless, the interplay between serotonin, dopamine, and cortisol in behavioral responses to threatening and stressful stimuli in zebrafish (see Soares, Gerlai, & Maximino, 2018, for a review) suggests a participation of uptake_2_. Here, we show that D-22 dose-dependently increases anxiety-like behavior in the zebrafish light/dark test (LDT). These results suggest that uptake_2_ is present in this species, and that it functions as a mediator of stress and defensive behavior.

This manuscript is a complete report of all the studies performed to test the hypothesis of a dosedependent effect of D-22 on anxiety-like behavior. We report all data exclusions (if any), all manipulations, and all measures in the study.

## 2. Materials and methods

### 2.1. Animals and housing

A total of 100 animals were used. Animals were bought from a commercial vendor and arrived in the laboratory with an approximate age of 3 months (standard length = 13.2 ± 1.4 mm), and were quarantined for two weeks; the experiment began when animals had an approximate age of 4 months (standard length = 23.0 ± 3.2 mm). Animals were kept in mixed-sex tanks during acclimation, with an approximate ratio of 50-50 males to females (confirmed by body morphology). Adult zebrafish from the wildtype strain (longfin phenotype) were used in the experiments. Outbred populations were used for increased genetic variability, thus decreasing the effects of random genetic drift which could lead to the development of uniquely heritable traits (Parra *et al*. 2009; Speedie and Gerlai 2008). Thus, the animals used in the experiments were expected to better represent the natural populations in the wild. The breeder was licensed for aquaculture under Ibama’s (Instituto Brasileiro do Meio Ambiente e dos Recursos Naturais Renováveis) Resolution 95/1993. Animals were group-housed in 40 L tanks, with a maximum density of 25 fish per tank, for at least 2 weeks before experiments begun. Tanks were filled with non-chlorinated water at room temperature (28 °C) and a pH of 7.0-8.0. Lighting was provided by fluorescent lamps in a cycle of 14-10 hours (LD), according to standards of care for zebrafish (Lawrence 2007). Water quality parameters were as follows: pH 7.0-8.0; hardness 100-150 mg/L CaCO3; dissolved oxygen 7.5-8.0 mg/L; ammonia and nitrite < 0.001 ppm. All manipulations minimized their potential suffering of animals, and followed Brazilian legislation (Conselho Nacional de Controle de Experimentação Animal - CONCEA 2017). Animals were used for only one experiment and in a single behavioural test, to reduce interference from apparatus exposure. Experiments were approved by UEPA’s IACUC under protocol 06/18.

### 2.2. Sample size calculation and exclusion criteria

Sample sizes were calculated based on a power analysis, using the effects of fluoxetine on the light/ dark test (Maximino *et al*. 2013) as estimates of effect sizes. Using an effect size of 0.7, a significance level of 0.005, and a power of 90%, a sample size of 11 animals per group was calculated. Final sample sizes were 20 animals/group. Animals were excluded if they displayed signals of overt ataxia (swimming on a side, swimming upside-down, vertical swimming; Demin et al., 2017) during the exposure period. Outliers were detected using an *a priori* rule based on median absolute deviation (MAD) of time on white (the main endpoint of the LDT), with values above or below 3 MADs being removed (Leys *et al*. 2013).

### 2.3. Drug treatments

Zebrafish were randomly drawn from the holding tank immediately before injection and assigned to four independent groups *(n* = 20/group). Animals were injected with vehicle (Cortland’s salt solution) D-22 (0.01, 0.1, 1, or 10 mg/kg). The injection volume was 1 μL/0.1 g b.w. (Kinkel et al. 2010). The order with which groups were tested was randomized via generation of random numbers using the randomization tool in http://www.randomization.com/. Experimenters were blind to treatment by using coded vials for drugs. The data analyst was blinded to phenotype by using coding to reflect treatments in the resulting datasets; after analysis, data was unblinded.

### 2.4. Light/dark test

The light/dark preference (scototaxis) assay was performed as described elsewhere (Maximino, 2018 [https://10.17504/protocols.io.srfed3n]; Maximino et al., 2010b). Briefly, 30 min after injection animals were transferred individually to the central compartment of a black/white tank (15 cm height X 10 cm width X 45 cm length) for a 3-min acclimation period, after which, the doors which delimit this compartment were removed and the animal was allowed to freely explore the apparatus for 15 min. While the whole experimental tank was illuminated from above by a homogeneous light source, due to the reflectivity of the apparatus walls and floor average illumination (measured just above the water line) above the black compartment was 225 ± 64.2 (mean ± S.D.) lux, while in the white compartment it was 307 ± 96.7 lux. The following variables were recorded:

1. *Time spent on the white compartment:* the time spent in the white half of the tank (*s*);
2. *Transitions to white:* the number of entries in the white compartment made by the animal throughout the session;
3. *Entry duration:* the average duration of an entry (time on white / transitions);
4. *Erratic swimming:* defined as the number of zig-zag, fast, and unpredictable swimming behavior of short duration;
5. *Freezing:* the duration of freezing events (*s*), defined as complete cessation of movements with the exception of eye and operculum movements;
6. *Thigmotaxis:* the duration of thigmotaxic events (*s*), defined as swimming in a distance of 2 cm or less from the white compartment’s walls;
7. *Frequency of risk assessment:* defined as a fast (<1 s) entry in the white compartment followed by re-entry in the black compartment, or as a partial entry in the white compartment (i.e., the pectoral fin does not cross the midline);

A digital video camera (Samsung ES68, Carl Zeiss lens) was installed above the apparatus to record the behavioral activity of the zebrafish. Two independent observers, blinded to treatment, manually measured the behavioral variables using X-Plo-Rat 2005 (https://github.com/lanec-unifesspa/x-plo-rat). Inter-observer reliability was at least > 0.95.

### 2.5. Data analysis

Drug effects were assessed using asymptotic general independence tests, using the R package ‘coin’ (Hothorn *et al*. 2006). Post-hoc analysis was made using pairwise permutation tests with correction for the false discovery rate. Data were presented as Cumming estimation plots, with Hedges’ *g* used to estimate effect sizes. Cumming estimates were made using 5000 bootstrap, and confidence intervals were bias-corrected and accelerated.

## 3. Results

4 animals were removed from analysis in the highest dose group due to overt ataxia, and 1 animal was removed from the 1 mg/kg group for the same reason. 5 animals (2 in the 0.1 mg/kg group, 1 in the 1 mg/kg group, and 1 in the 10 mg/kg group) were detected as outliers and removed from further analysis. A dose-dependent decrease in time on white was found (maxT = −3.773, *p* = 0.0007; Figure 1B); significant effects were found for 0.1-10 mg/kg. Likewise, dose-dependent decreases were found for transitions to white (maxT = 4.0277, *p* = 0.0003; Figure 1C); significant effects were found for all doses, except 0.1 mg/kg. No significant effects were found for entry duration (maxT = 1.8191, *p* = 0.2779; Figure 1D). An inverted-U-shaped response was found for risk assessment (maxT = 4.6248, *p* = 0.019; Figure 1E), with 0.1, and 1 mg/kg increasing risk assessment, and 0.01 mg/kg having no effect; the effect of 10 mg/kg was smaller than the other effects. A main effect of dose was found in erratic swimming (maxT = 3.106, *p* = 0.0091; Figure 1F), but *post-hoc* comparisons failed to detect differences. No effects were found for thigmotaxis (maxT = 2.1474, *p* = 0.1396; Figure 1G) or freezing (maxT = 2.0629, *p* = 0.1688; Figure 1H).

**Figure 1.**
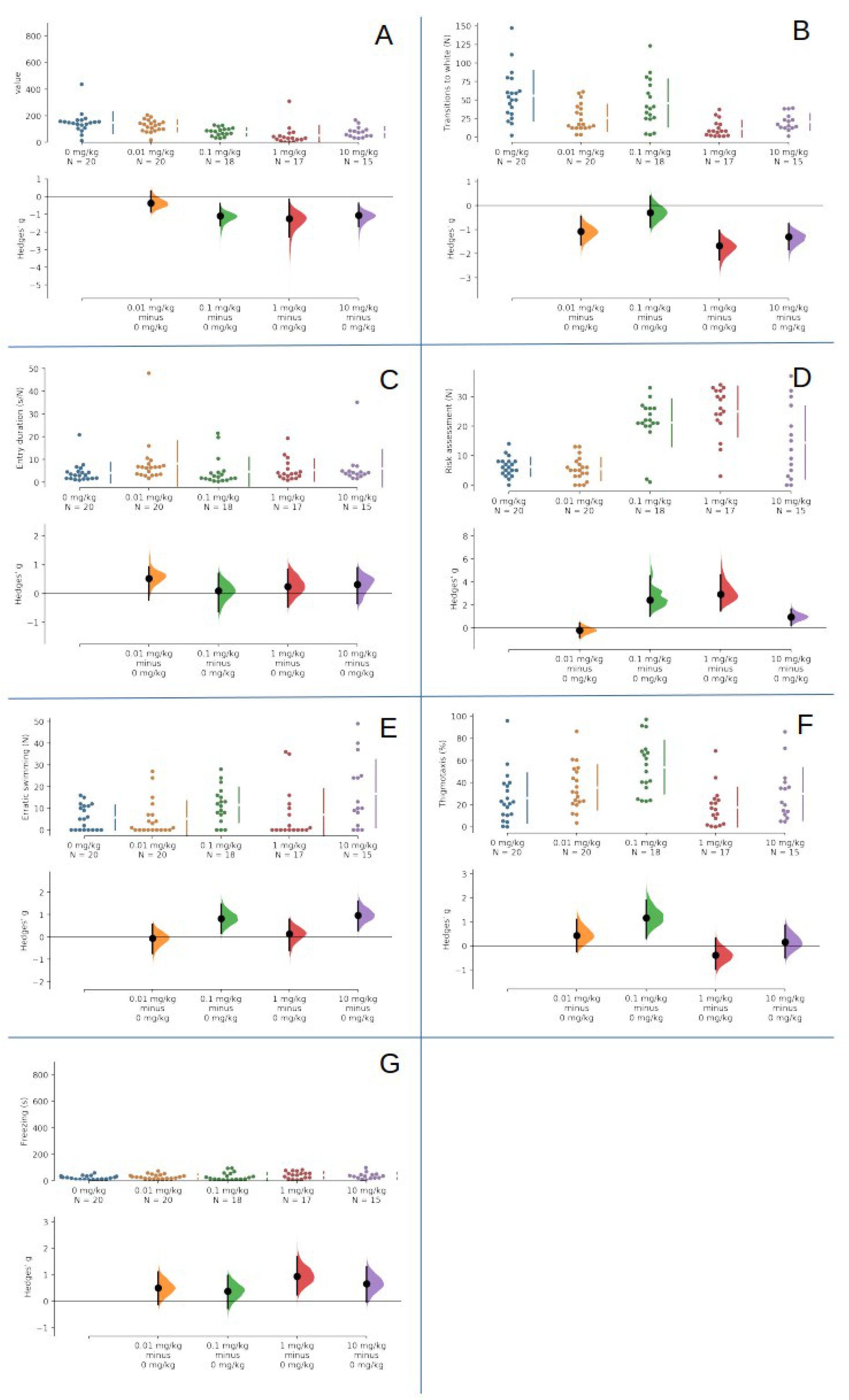
Decynium-22 increases anxiety-like behavior in the zebrafish LDT. (A) Scototaxis (time spent in the white compartment); (B) Transitions to the white compartment. (C) Duration of entries in the white compartment. (D) Risk assessment. (E) Erratic swimming. (F) Thigmotaxis. (G) Freezing duration. The Hedges′ *g* for 4 comparisons against the shared control 0 mg/kg are shown in the above Cumming estimation plots. The raw data is plotted on the upper axes. On the lower axes, mean differences are plotted as bootstrap sampling distributions. Each mean difference is depicted as a dot. Each 95% confidence interval is indicated by the ends of the vertical error bars. 5000 bootstrap samples were taken; the confidence interval is bias-corrected and accelerated. Letters indicate results from post-hoc tests; different letters indicate statistically significant differences (p < 0.05).

## 4. Discussion

The present experiment showed evidence that D-22, an uptake_2_ inhibitor, dose-dependently increased anxiety-like behavior in the LDT in unstressed zebrafish. Dose-dependent effects were found for time on white (scototaxis) and risk assessment, with the latter suggesting better effects at intermediate doses (0.1 and 1 mg/kg). No effects were observed in other variables (freezing and erratic swimming, thigmotaxis).

The LDT has been proposed as a screening test for anxiolytic-like and anxiogenic-like effects of treatments in adult zebrafish (Maximino *et al*. 2010a). The test shows good predictive validity, being sensitive to agents that act at different targets (Kysil *et al*. 2017). The main endpoint of this test, scototaxis, is sensitive to anxiolytic-like and anxiogenic-like effects, and represents an “avoidance” dimension of behavior in the LDT, while risk assessment clusters in a different group and represents a more “cognitive” aspect of anxiety-like behavior (Maximino *et al*. 2014). Moreover, exposure to the LDT induces a cortisol response in unstressed animals (Kysil *et al*. 2017), suggesting that the conflict that is induced in the test is mildly stressful.

Behavioral effects of D-22 have been described in rodents; while by itself D-22 (0.01-0.32 mg/kg) was not able to change immobility time in the tail suspension test in mice, a screen for antidepressant-like effects, it produced a synergistic effect with fluvoxamine (Horton *et al*. 2013). Species- and strain-specific effects can be responsible for this lack of effect of D-22, as this drug (0.001-0.01 mg/kg) reduced immobility time in the forced swim test (another screen for antidepressant-like effects) in Wistar-Kyoto, but not Long Evans, rats (Marcinkiewcz and Devine 2015). Although these effects are usually attributed to effects on serotonin clearance (Daws 2009b), it is not possible to discard an effect on norepineprhine.

D-22 blocks the uptake_2_ monoamine transport system (Daws 2009b). Due to its low-affinity, high-capacity character, transporters in the uptake_2_ system (OCT and PMAT) are “promiscuous”, participating in the elimination of most monoamines from synaptic and extrasynaptic sites (Hagan *et al*. 2011). Importantly, uptake_2_ may represent a link between acute stress and monoaminergic neurotransmission (Maximino 2012), as these transporters are blocked by glucocorticoids (Hill *et al*. 2011). While currently it is not known whether the effects reported in this experiment are due to serotonin, norepinephrine, dopamine, or histamine, there is some evidence for anxiety-like behavior in zebrafish being increased by serotonin (Herculano and Maximino 2014) and catecholamines (Idalencio *et al*. 2017a).

Overall, these results suggest that uptake_2_ is present in zebrafish, and that it functions as a mediator of stress and defensive behavior. These results point to novel avenues of investigation in the stress-monoamine interaction in anxiety, stress, and defensive behavior. Further studies are needed to better understand the mechanisms by which D-22 produces its behavioral effects.

## Significance statement

Uptake_2_ is a low-affinity, high-capacity transport system that contributes to the clearance of extraneuronal monoamines (mainly norepinephrine, serotonin, and dopamine) and is sensitive to glucocorticoids, therefore representing a putative mechanism of glucocorticoid-monoamine interaction. Since both monoamines and glucocorticoids have been implicated as mediators of stress-induced behavioral adjustments, this interaction can of relevant to understand the mechanisms through which stress influences neurochemical and behavioral responses. Here we report that, in zebrafish, the uptake_2_ inhibitor decynium-22 increases dark preference and risk assessment in the light/dark test, an assay for anxiety-like behavior. Thus, uptake_2_ appears to act as a modulator of defensive behavior, and its inhibition by, e.g., glucocorticoids could represent a mechanism through which stress produces fast neurobehavioral adjustments in vertebrates.

## Authors’ contributions

*Caio Maximino:* Conceptualization, Formal analysis, Funding acquisition, Methodology, Investigation, Data curation, Project Administration, Resources, Software, Supervision, Validation, Visualization, Writing – original draft.

## Data accessibility

Data and analysis scripts for this work can be found at a GitHub repository (https://github.com/lanec-unifesspa/decynium22).

